# A 20+ Ma old enamel proteome from Canada’s High Arctic reveals diversification of Rhinocerotidae in the middle Eocene-Oligocene

**DOI:** 10.1101/2024.06.07.597871

**Authors:** Ryan S. Paterson, Meaghan Mackie, Alessio Capobianco, Nicola S. Heckeberg, Danielle Fraser, Fazeelah Munir, Ioannis Patramanis, Jazmín Ramos-Madrigal, Shanlin Liu, Abigail D. Ramsøe, Marc R. Dickinson, Chloë Baldreki, Marisa Gilbert, Raffaele Sardella, Luca Bellucci, Gabriele Scorrano, Fernando Racimo, Eske Willerslev, Kirsty E.H. Penkman, Jesper V. Olsen, Ross D.E. MacPhee, Natalia Rybczynski, Sebastian Höhna, Enrico Cappellini

## Abstract

In the past decade, ancient protein sequences have emerged as a valuable source of data for deep-time phylogenetic inference. Still, the recovery of protein sequences providing novel phylogenetic insights does not exceed 3.7 Ma (Pliocene). Here, we push this boundary back to 21-24 Ma (early Miocene), by retrieving enamel protein sequences of an early-diverging rhinocerotid (*Epiaceratherium* sp. - CMNF-59632) from the Canadian High Arctic. We recover partial sequences of seven enamel proteins (AHSG, ALB, AMBN, AMELX, AMTN, ENAM, MMP20) and over 1000 peptide-spectrum matches, spanning over at least 251 amino acids. Authentic endogeneity of these sequences is supported by indicators of protein damage, including several spontaneous and irreversible post-translational modifications accumulated during prolonged diagenesis and reaching near-complete occupancy at many sites. Bayesian tip-dating, across 15 extant and extinct perissodactyl taxa, places the divergence time of CMNF-59632 in the middle Eocene-Oligocene, and identifies a later divergence time for Elasmotheriinae in the Oligocene. The finding weakens alternative models suggesting a deep basal split between Elasmotheriinae and Rhinocerotinae. This divergence time of CMNF-59632 coincides with a phase of high diversification of rhinocerotids, and supports a Eurasian origin of this clade in the late Eocene or Oligocene. The findings are consistent with previous hypotheses on the origin of the enigmatic fauna of the Haughton crater, which, in spite of their considerable degree of endemism, also display similarity to distant Eurasian faunas. Our findings demonstrate the potential of palaeoproteomics in obtaining phylogenetic information from a specimen that is ten times older than any sample from which endogenous DNA has been obtained.

---

Phylogenetic placement of deep-time (>1 Ma) fossils has typically relied on morphological observations, as the recovery of sufficiently extensive genetic evidence has not been thought to be possible beyond the Pleistocene^1^. While ancient DNA (aDNA) sequences are a valuable source of data for inferring phylogenies and population dynamics in the late Pleistocene^2–5^, the oldest authentic aDNA from macrofossils has been extracted from Arctic-situated specimens dated to no more than 1.2 Ma^6^. In contrast, palaeoproteomic data have been recovered from late Miocene, Pliocene and early Pleistocene fossils, even in localities that are warm, humid, and/or at low latitudes^7,8^. Although protein sequences from the early Pleistocene have been successfully used to infer the phylogenetic placement of various fossil mammals^9–11^, the precise limit of proteomic survival has not yet been systematically characterised^12,13^. Currently, the oldest confirmed palaeoproteomic data successfully used to infer sub-ordinal taxonomic relationships derive from bone collagen of camelids from the 3.7 Ma Fyles Leaf Bed site of Canada’s High Arctic^14,15^. This illustrates how our understanding of evolutionary relationships is currently limited by the preservation of biomolecules from extinct species.

Rhinocerotidae is a family that includes only five extant species, but a wide diversity of fossil members^16,17^. It remains debated as to where and when the radiation of this group occurred^18^. For most of the last two decades, the group was defined by a deep ‘basal split’ between two clades - Rhinocerotinae and Elasmotheriinae - prior to bursts of rhinocerotid diversification in the late Eocene^19–23^. This paradigm contrasts earlier hypotheses of a close relationship between two extinct rhinocerotids that survived into the late Pleistocene – the Siberian unicorn (Elasmotheriinae, *Elasmotherium sibiricum*) and the woolly rhino (Rhinocerotinae, *Coelodonta antiquitatis*)^24^. Recently, the sequenced genomes of *Coelodont*a and *Elasmotherium^25^*were used to confirm hypotheses based on morphological data that suggest they have distinct phylogenetic affinities^19^, but also allowed for the recognition of a relatively-young split between these two groups during the late Eocene (36 Ma). This suggests that the deep-divergence hypothesis based on the morphological analysis of fossils is not supported by molecular evidence. However, the lack of available genetic sequence data from other early-diverging rhinocerotid lineages makes it difficult to assess the timing of the Rhinocerotinae-Elasmotheriinae split in relation to other radiations that occurred within the group. For these reasons, the ancient radiations of the group still remain obscured.

To investigate the timing of Rhinocerotidae divergence and the potential for evolutionarily informative protein sequences to persist in deep-time, we targeted vertebrate dental enamel deriving from the Haughton crater (75°N, Nunavut) in Canada’s High Arctic (Figure 1). The Haughton crater is an impact structure with its stratigraphy including post-impact fossiliferous lacustrine sediments dated to 21-24 Ma^26^. While their geological age is advanced, fossils from these sediments are found in a polar landscape, currently characterised by permafrost. This creates a temperature regime favourable for biomolecular preservation, sparing these fossils from the harshest effects of diagenesis. Accordingly, specimens from this site serve as promising candidates for biomolecular preservation in deep-time.

**Figure 1).**
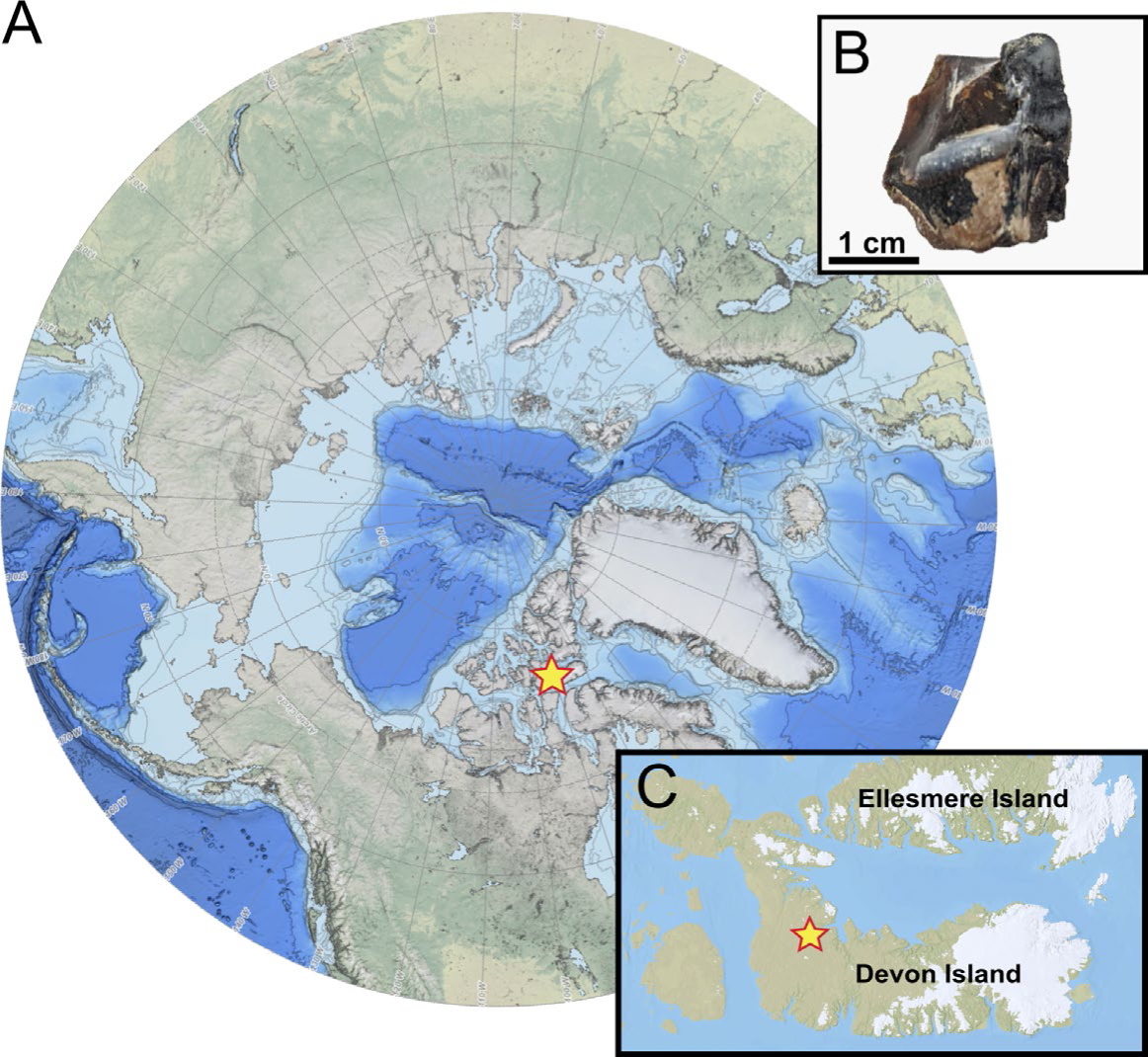
The high latitude Haughton crater on Devon Island has produced a highly endemic vertebrate fauna. A) Location of Devon Island within the circumpolar North. B) Anterolingual view of specimen CMNF-59632 after destructive palaeoproteomic analysis. C) Location of Haughton crater on Devon Island.

The digestion-free palaeoproteomic workflow^9,10^ applied to an early Miocene rhinocerotid (*Epiaceratherium* sp.) specimen of dental enamel^27^ from the Haughton Formation (21.8 Ma) allowed for the recovery of an enamel proteome covering 1163 confident peptide-spectrum matches (PSMs), at least seven proteins (AHSG, ALB, AMBN, AMELX, AMTN, ENAM, MMP20), and spanning at least 251 amino acids (Figure 2a) (Ext. Figure 1a). The enamel proteome of CMNF-59632 currently represents both the oldest mammalian skeletal proteome currently reported, confirming the predicted deep-time persistence of ancient mammalian proteins from high latitudes^8,10^, and the first biomolecular characterisation of the extinct genus *Epiaceratherium*. While the survival of a relatively rich enamel proteome from such ancient deposits is surprising, the age of the specimen belies its excellent state of preservation.

**Figure 2).**
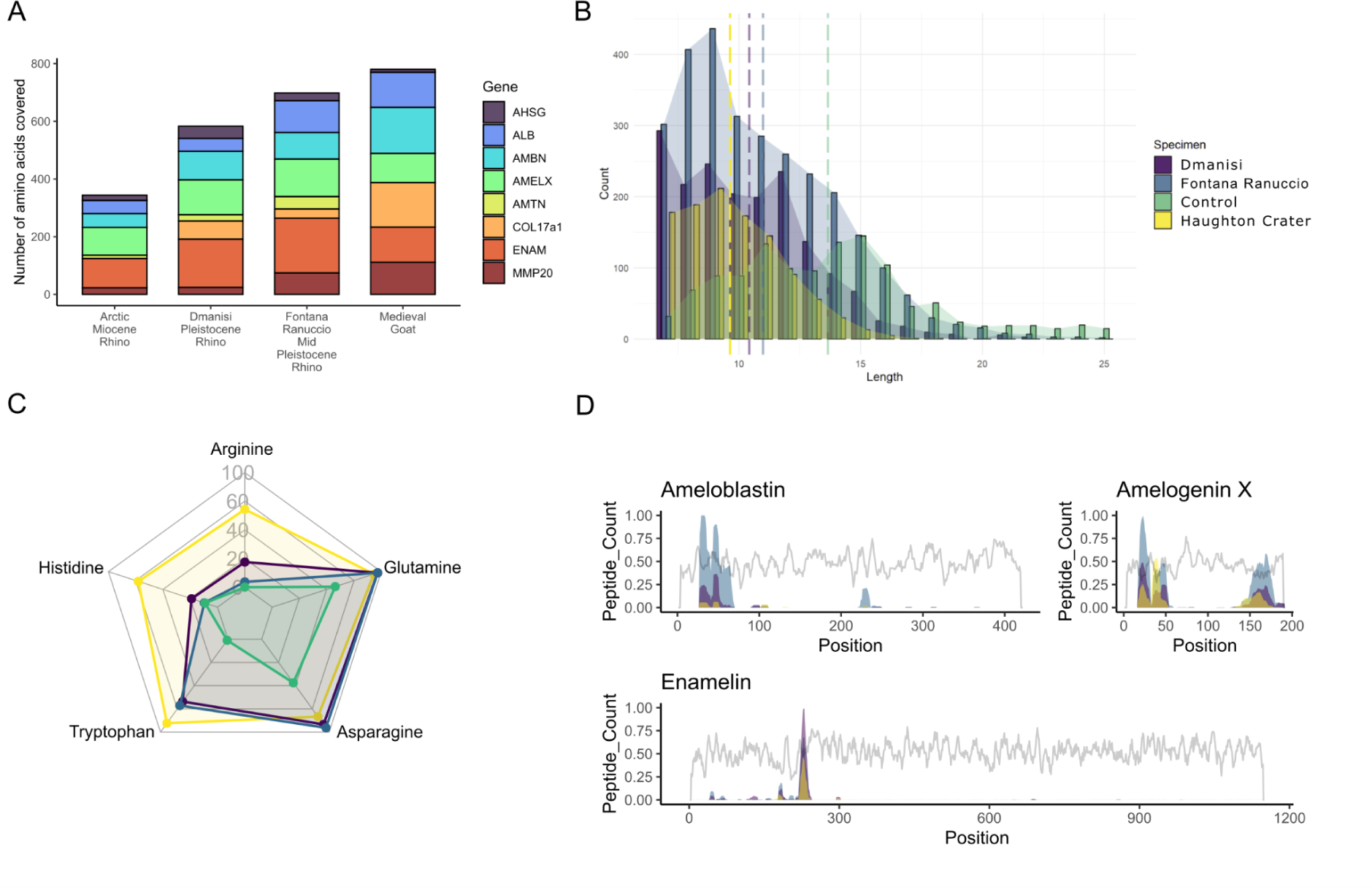
Proteome preservation in the enamel specimen of the early Miocene rhinocerotid (CMNF-59632). Preservation is compared to enamel proteomes from an early Pleistocene (1.77 Ma) *Stephanorhinus* (DM.5/157), a middle Pleistocene (0.4 Ma) *Stephanorhinus* (CGG 1_023342) and a medieval ovicaprine (Control)^9^. All plots exclude contaminants and reverse hits. A) Amino acid sequence coverage for each identified protein; B) peptide length distributions, dashed bars represent average peptide length for each specimen; C) proportion of a selected sample of amino acids that are often modified in ancient enamel proteomes. Results derive from PTM-specific searches described in methods. ‘Arginine’ includes arginine-to-ornithine conversion, ‘Glutamine’ includes glutamine deamidation, ‘Asparagine’ includes asparagine deamidation, ‘Tryptophan’ includes advanced tryptophan oxidation to kynurenine, oxylactone, and tryptophandione, ‘Histidine’ includes oxidation and dioxidation of histidine, histidine conversion to hydroxyglutamate. D) Sequence coverage plots for the three most abundant enamel matrix proteins (AMBN, AMELX, ENAM), recording number of PSMs (coloured areas) and mutability (grey line).

To better appreciate the preservation state of the Haughton Crater enamel proteome, we compared it to those of two other rhinocerotids, namely the early Pleistocene *Stephanorhinus* from the site of Dmanisi (Georgia), dated at 1.77 Ma^9^, and a middle Pleistocene *Stephanorhinus* (∼0.4 Ma) from the site of Fontana Ranuccio (Italy). While the set of proteins retrieved from the CMNF-59632 enamel specimen is similar to that of the other two Pleistocene rhinocerotids used for comparison (Figure 2a), fewer peptides and a shorter reconstructed amino acid sequence were recovered from the Arctic specimen. In addition to proteins previously found in deep-time enamel samples (AHSG, AMBN, AMELX, AMTN, ENAM, MMP20), serum albumin was also found in each rhinocerotid sample. Previously recovered from other fossil enamel specimens^11^, serum albumin is phylogenetically more informative than enamel-specific proteins due to its higher amino acid sequence variability^28^. In the present analysis, while some albumin peptides are removed during initial data filtering (as they are identical to peptide sequences from bovine serum albumin, a potential laboratory contaminant), several others do not match contaminants. The vast majority of the spectra that confidently support their identification show post-translational modifications (PTMs) that derive from prolonged diagenesis, which supports their authenticity.

As expected, diagenetic modifications and PTMs are extensive in the enamel proteome of CMNF-59632 (Figure 2b). Average peptide lengths are similar, though slightly shorter than those of the Dmanisi early Pleistocene specimen, and further reduced in comparison to the Fontana Ranuccio middle Pleistocene *Stephanorhinus*, indicating a greater degree of peptide bond hydrolysis (Figure 2b). We also observe high deamidation rates in CMNF-59632, though no more so than in the Pleistocene rhinocerotids (Ext. Figure 2). While high deamidation rates can be useful for confirming proteome authenticity, they can be highly variable within samples^29–31^, and can plateau relatively-quickly in fossil proteomes, reducing their utility in characterising degradation patterns in deep-time (Figure 2c). Instead, we identify a suite of informative spontaneous PTMs indicative of advanced diagenesis that are observed at a higher rate in the Arctic Miocene rhinocerotid, providing support for their utility as markers of advanced diagenesis and authenticity in deep-time (Figure 2c)^9^. These include arginine to ornithine conversion (Figure 2c) and advanced forms of tryptophan (Ext. Figure 2b) and histidine oxidation (Ext. Figure 1c). Intra-crystalline protein decomposition analysis further confirms the advanced degradation state of CMNF-59632. The concentration of free and total hydrolysable amino acids (FAA and THAA, respectively) is around half of those in the early Pleistocene *Stephanorhinus* sample from Dmanisi (Extended Figure 2a), and the percentage of FAA in CMNF-59632 (∼75%) is higher than in the Pleistocene rhino from Dmanisi (∼50%) (Ext. Figure 3b), supporting increased peptide bond hydrolysis. Furthermore, the racemisation values for CMNF-59632 fall along the expected FAA vs THAA trends for both fossil enamel, and the experimentally-heated samples (Ext. Figure 4), confirming the closed system behaviour of CMNF-59632 enamel amino acids, and supporting the endogeneity of the peptides retrieved.

At least 10 single amino acid polymorphisms (SAPs) support the placement of CMNF-59632 within Rhinocerotidae. A smaller number (2+) of SAPs are shared between CMNF-59632 and other perissodactyls, to the exclusion of later-diverging rhinocerotids. No novel variants are uncovered in CMNF-59632, as the aforementioned SAPs represent character states retained from ancestors within Perissodactyla and Mammalia more broadly. The identification of these SAPs is supported by several unique PSMs displaying almost complete ion series (e.g., Ext. Figure 5).

Peptide sequences recovered from CMNF-59632 derive from similar sequence regions to those previously identified in the Dmanisi Pleistocene *Stephanorhinus* proteome (Figure 2d), particularly for the three most abundant enamel matrix proteins (EMPs). ENAM and AMBN present broadly similar sequence coverage patterns in both specimens, though with fewer PSMs covering most positions in the Miocene sample. AMELX, the most abundant EMP, is instead covered by a similar number of PSMs in both the Miocene and Pleistocene samples. The depth of coverage is also similar for the most abundantly-covered AMELX sequences, including those spanning the deletion observed in the Leucine-Rich Amelogenin Peptide (LRAP)^9,32^.

Regardless of the mechanisms behind preferential mass spectrometric and data analysis identification of specific sequence regions, biases favouring the recovery^7^ and identification^33^ of conserved peptide sequences can ultimately lead to underestimates of divergence times in taxa represented by empirically-derived protein sequences. To accurately estimate the phylogenetic position of CMNF-59632 and estimate divergence times within the group, we completed a phylogenetic analysis of a suite of extinct and extant perissodactyls. In addition to the perissodactyl taxa used in Cappellini et al. (2019)^9^, we incorporated whole-genome sequence data to predict enamel protein sequences from the Siberian unicorn (*Elasmotherium sibiricum*) and a pair of extant tapirs (*Tapirus terrestris* and *Tapirus indicus*).

The time-calibrated phylogenetic analysis of enamel protein sequences under a Fossilised Birth Death (FBD) model infers CMNF-59632 as the earliest diverging rhinocerotid in the analysis, with *Elasmotherium sibiricum* being more closely related to Rhinocerotina (crown rhinoceroses) than to CMNF-59632 (Figure 4). This phylogenetic hypothesis is also supported by Fraser et al. (2024) in a total-evidence analysis. Additionally, our FBD analysis resolves the early Pleistocene *Stephanorhinus* from Dmanisi as a sampled ancestor of the middle Pleistocene *Stephanorhinus* from Fontana Ranuccio. Divergence time estimates place the split between CMNF-59632 and all other rhinocerotids during the middle Eocene–Oligocene (around 41–25 Ma). The divergence between *Elasmotherium sibiricum* and Rhinocerotina is reconstructed to have likely occurred in the Oligocene (around 34–22 Ma), which is younger than previous molecular clock estimates^25^.

**Figure 3).**
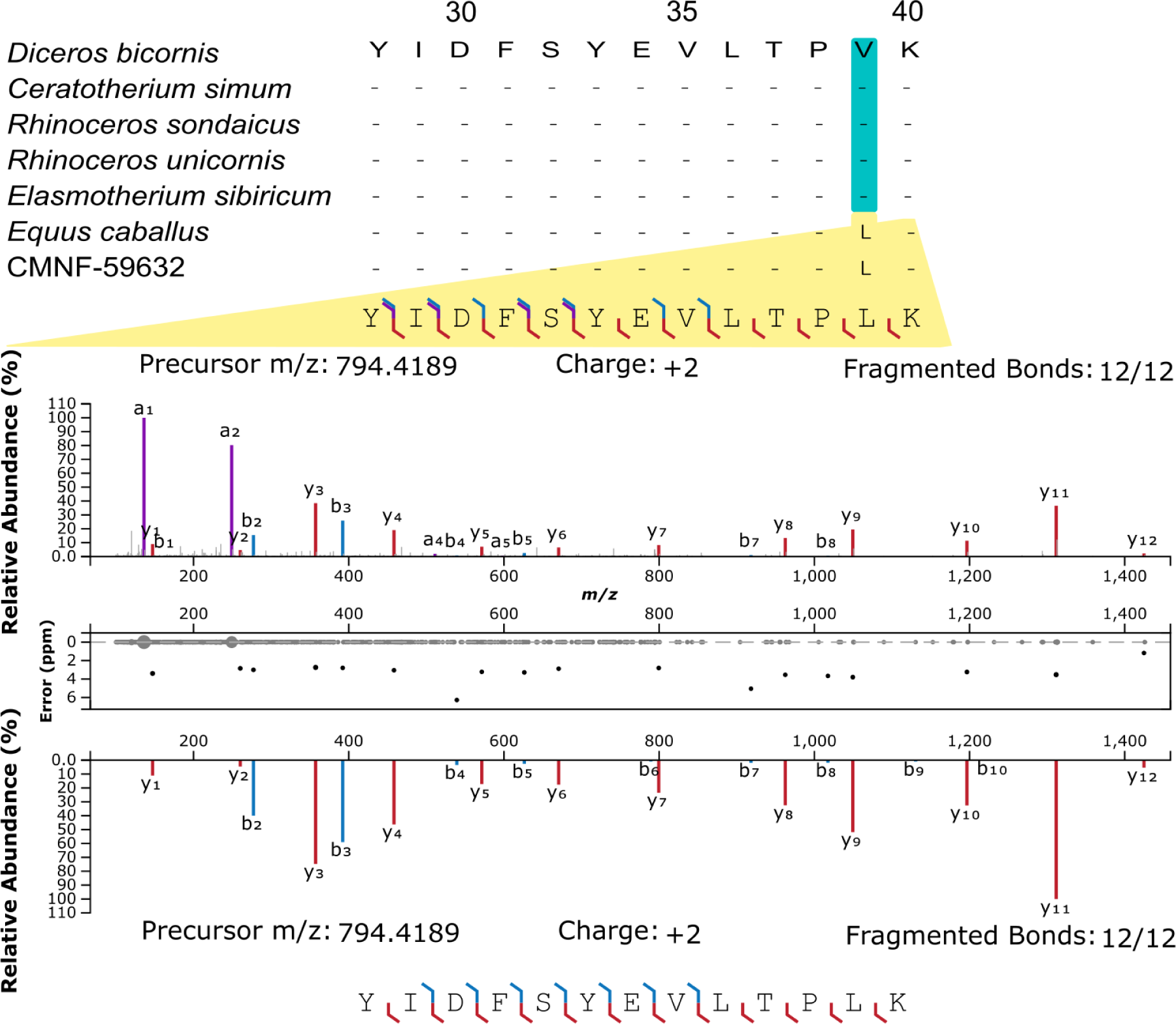
Abridged alignment and mirror plots of a phylogenetically-informative single amino acid polymorphism (SAP) at AMELX-39. The top spectrum is experimentally-derived, while the bottom one is predicted using the ‘Original mode’ with the Prosit tool, available online via the Universal Spectrum Explorer^83^. This spectrum is the highest scoring peptide-spectrum match (with Andromeda) for AMELX sequence positions spanning the most abundantly-covered SAP differentiating between CMNF-59632, and all other rhinocerotids for which sequences are available.

**Figure 4).**
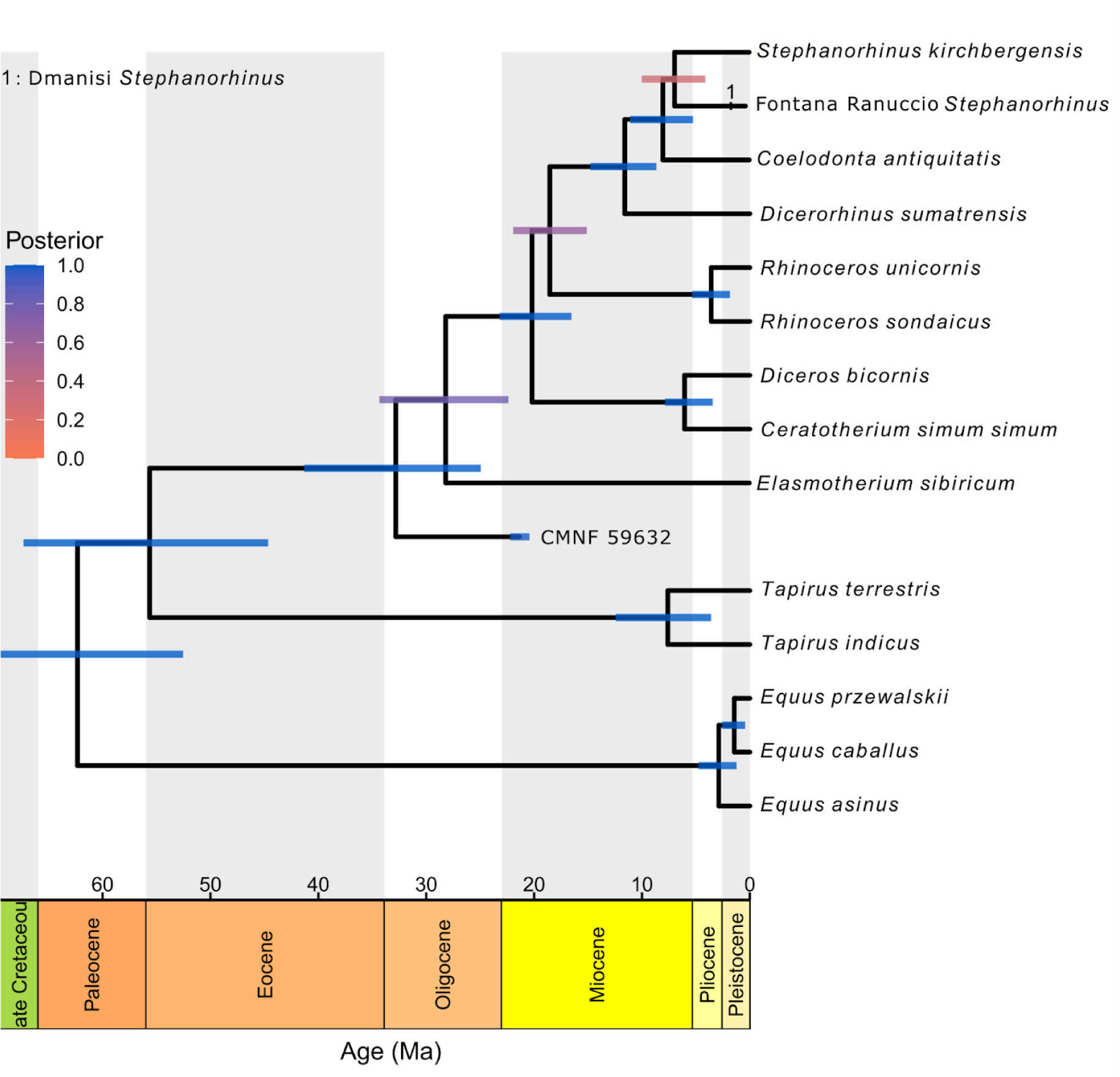
Time-calibrated phylogeny of Rhinocerotidae enamel proteomes. The maximum *a posteriori* (MAP) tree was produced using RevBayes v.1.2.1 ^70^; https://revbayes.github.io, with a Fossilized Birth Death (FBD) model. Coloured bars at nodes represent 95% height posterior density (HPD) age interval estimates. Specimen CMNF-59632 represents the early Miocene rhinocerotid from the Haughton Crater.

The late Eocene and the early Oligocene represent dynamic periods in the evolution of rhinocerotids, particularly in North America. After appearing in the middle Eocene (37-34 Ma)^34^, North American rhinocerotids diversify during the late Eocene, evolving a variety of body sizes and ecologies as several new clades arise, before rhinocerotid diversity experiences a significant drop in the early Oligocene (34-32 Ma)^35^. During this timeframe, other early-diverging lineages are also appearing in Asia^18,36,37^, and eventually spreading as far as Western Europe^18^. Morphologically, the Haughton crater rhinocerotid shares closer affinities with these early-diverging lineages from Eurasia^38^, particularly those within the genus *Epiaceratherium^27^*. Similarly, some other vertebrates within the highly-endemic fauna of the Haughton Formation have their closest relatives in Eurasia. These include the transitional pinniped *Puijila darwini*, sister to the Oligocene *Potamotherium* of Europe^39^, and the swan-like anatid, a group which is otherwise restricted to the Oligocene and Miocene of Europe^38^. Overall, these patterns, in conjunction with the recovered divergence times, suggest the Haughton crater rhinocerotid represents a migrant from eastern Asia or western Europe, derived from one of the early-diverging lineages that arose in the late Eocene or early Oligocene of East Asia.

We provide molecular evidence that this lineage falls outside of Rhinocerotinae, as it diverges before the Rhinocerotinae-Elasmotheriinae split. We also reject a deep-divergence (basal split) between Elasmotheriinae and Rhinocerotinae^19–23^ and find moderate support for their branching event after the divergence of *Epiaceratherium*. Our analysis disagrees with that of Kosintsev et al. (2019)^23^, who find a deep divergence for Elasmotheriinae (47.3 Ma), and an early divergence for Rhinocerotinae (almost 30.8 Ma). The later divergence times for these nodes in our analysis are in spite of equivalently old ages for crown Ceratomorpha (earliest Eocene). Among other timetrees, our dates are generally most consistent with those of Liu et al. (2021)^25^. Our recovered topologies are also broadly similar to trees derived from the morphology-based phylogenetic analyses of Tissier et al. (2020)^18^ and Lu et al. (2023)^40^, identifying Elasmotheriinae and Rhinocerotinae as deeply-nested within Rhinocerotidae. Discrepancies between the genomic ^25^ and proteomic trees arise likely due to different calibration points. The more ancient age of Elasmotheriinae in the analysis of Liu et al. (2021)^25^ is constrained by a high minimum bound for the Elasmotheriinae-Rhinocerotinae split (35 Ma). However, this date is based on the earliest age of *Epiaceratherium naduongense* and its allocation to Rhinocerotinae. Assuming monophyly of *Epiaceratherium*, the present proteomic evidence refutes the assignment of this genus to Rhinocerotinae, as it falls as earlier-diverging than Elasmotheriinae without such topological constraints in our phylogenetic analysis.

In sum, these findings highlight the importance of integrating palaeoproteomic sequence data into phylogenetic analyses to infer topologies and estimate divergence times. Ancient proteomic sequence data allows for robustly-supported timetrees, and can serve to develop phylogenetic frameworks in deep-time, particularly from specimens too old to preserve ancient DNA. For example, the present data allows for firm placement of the Haughton Crater rhinocerotid outside of Rhinocerotina, and likely outside the Elasmotheriinae-Rhinocerotinae clade, a fact which has significant implications for both morphological and molecular studies integrating fossil calibration times from the fossil record. Furthermore, we demonstrate the deep-time survival of a rich set of peptides derived from proteins present in mammalian enamel, well beyond the previously known limits of survival. This work illustrates the power of palaeoproteomics in elucidating phylogeny and taxonomy of extinct vertebrates in deep-time. These findings should encourage further vertebrate palaeontological fieldwork in the High Arctic, and other cold-temperature sites with taphonomic conditions favourable to biomolecular preservation.

## Materials and Methods

### Site and Specimen

Located within the Haughton impact crater (75°N, Nunavut, Canada), the Haughton Formation comprises the remnants of a large, post-impact lacustrine deposit, dated to the early Miocene. Previous dating estimates, using fission-track and ^40^Ar–^39^Ar furnace step-heating dating, identified an age of 24-21 Ma^26,41^. An early Miocene age has also been corroborated by (U-Th)/He thermochronology^42^. While older age estimates between 30-40 Ma have also been suggested^43–45^, there have been no age estimates younger than the early Miocene. Therefore, we conservatively use the younger early Miocene age estimates in our analysis and interpretation.

The highly-endemic fauna of the Haughton formation consists of several vertebrate taxa, including a transitional pinniped^39^, a pair of salmoniform fishes, a swan-like anatid, a small artiodactyl, a leporid rabbit, a heterosocid shrew, and a well-preserved rhinocerotid^27,38^. While the megafloral assemblage is not particularly rich, the palynofloral assemblage is well-characterised, allowing for reconstruction of local climatic conditions. In the early Miocene, the Haughton crater lake and its surrounding environs experienced a significantly warmer annual temperature (8-12°C) than the present day^38,46^.

The specimen CMNF-59632 is a nearly complete rhinocerotid skeleton, including skull and dentition, uncovered 10.8 m above the base of the formation^27^. The present analysis focuses on a single tooth fragment from a lower left m1 (Figure 1b) that was already separated from the rest of its tooth row due the fragmenting effectings of cryoturbation. The dental specimen’s rhinocerotid affinities are further supported by its size and morphology, most notably the presence of vertical Hunter-Schreger bands on its enamel, a defining feature of rhinocerotids and found in few other mammals^47^. A single tusk fragment (left i2) derived from CMNF-59632 was also selected for proteomic extraction. Due to its thin enamel, only limited peptides were recovered from this tusk fragment, and the sample is thus excluded from further analysis and discussion.

### Proteomic extraction and LC-MS/MS

The laboratory workflow for the CMNF-59632 teeth and the Fontana Ranuccio *Stephanorhinus* tooth (for comparison) generally follows that of Cappellini et al. (2019) and Taurozzi et al. (2024)^9,48^. Using a sterilised drill, flakes of enamel were removed from the fragmentary teeth, with care taken to avoid sampling the dentine. The CMNF-59632 tooth enamel sample, 154 mg, was then ground to a fine powder, and demineralized overnight using 10% HPLC-grade trifluoroacetic acid (TFA) (Merck, Sigma-Aldrich). The CMNF-59632 tusk enamel sample, weighing 90 mg, was processed in the same way. The Fontana Ranuccio enamel sample was divided into three subsamples - FR2, FR3, and FR4 - weighing 202, 243, and 205 mg, respectively. They were similarly ground to a fine powder, and demineralized using 10% TFA (FR3, FR4) or 10% HCl (hydrochloric acid, FR2). For each sample, the demineralization step was repeated a second time to ensure complete demineralization. As enamel peptides are already hydrolysed *in vivo*, no enzymatic digestion was performed. Subsequently, peptides were collected and desalted on C-18 StageTips^49^ produced in-house. An extraction blank for each sample set was processed alongside the samples for every step, to control for contamination.

### Mass spectrometry

Stagetips were eluted with 30 μL of 40% acetonitrile (ACN) and 0.1% formic acid (FA) into a 96-well plate. To remove ACN and concentrate the samples, they were vacuum-centrifuged until approximately 3 μL of sample remained. Next, samples were resuspended in 6 μL of 5% ACN 0.1% formic acid (FA).

Liquid chromatography coupled with tandem mass spectrometry (LC-MS/MS) was used to analyse the samples, based on previously published protocols^9,50^. Samples were separated on a 15 cm column (75 μm inner diameter in-house laser pulled and packed with 1.9 μm C18 beads (Dr. Maisch, Germany) on an EASY-nLC 1200 (Proxeon, Odense, Denmark) connected to an Exploris 480 (CMNF-59632) or a Q-Exactive HF-X (Fontana Ranuccio *Stephanorhinus*) mass spectrometer (both from Thermo Scientific, Bremen, Germany), with an integrated column oven. 4 (CMNF-59632) or 5 (FR sd - 295 - *Stephanorhinus*) μL of sample were injected. 0.1% FA in milliQ water was used as buffer A and the peptides were separated with increasing buffer B (80% ACN 0.1% FA) with a 77 min gradient, increasing from 5% to 30% in 50 min, 30% to 45% in 10 min, 45% to 80% in 2 min, and maintained at 80% for 5 min before decreasing to 5% in 5 min, and finally held for 5 min at 5%. Flow rate was 250 nL/min. An integrated column oven was used to maintain the temperature at 40°C.

The two mass spectrometers were run with the same parameters except where specified, due to changes in running software. Spray voltage was set to 2 kV, the S-lens RF level was set to 40%, and the heated capillary was set to 275°C. Full scan mass spectra (MS1) were recorded at a resolution of 120,000 at m/z 200 over the m/z range 350-1400. The AGC target value was set to 300% / 3e6 (Exploris / HF-X) with a maximum injection time of 25 ms. HCD-generated product ions (MS2) were recorded in data-dependent top-10 mode and recorded at a resolution of 60,000. The maximum ion injection time was 118 / 108 ms (Exploris / HF-X), with an AGC target value of 200% / 2e5 (Exploris / HF-X). Normalised collision energy was set at 30 / 28% (Exploris / HF-X). The isolation window was set to 1.2 m/z with a dynamic exclusion of 20 s. A wash-blank, using 5% ACN, 0.5% TFA, was run between each sample and laboratory blank to limit cross-contamination.

### Database construction

The protein reference alignment used in Cappellini *et al*. (2019)^9^ was used as a starting point to construct a database for sequence reconstruction. Due to the vast evolutionary distance between CMNF-59632 and any extant taxa (>20 million years), a broader database was constructed to identify sequence variants that may be known in other mammals. To construct a broader database, we searched Uniprot and NCBI for each enamel protein, specifying the taxonomic grouping of ‘Theria’ to include all therian mammals. To supplement available sequences, additional sequences were manually extracted from available genomes, following the methodology from^51^.

To investigate the relationships at the base of Rhinocerotidae, protein sequences translated from *Elasmotherium sibiricum* genomic data^25^ were generated. To obtain the corresponding amino acid sequences, we firstly collapsed the Paired-End (PE) reads and masked the conflict bases as “N” using adapterRemoval^52^. Then, we mapped the collapsed reads against the reference genome of the white rhinoceros (GCF_000283155.1_CerSimSim1) using the BWA MEM function^53^ with the shorter split hits being abandoned. After that, we removed duplicates using an in-house Perl script following Liu et al. (2021)^25^. Finally, we extracted the gene sequences according to their locations on the reference genome.

The remaining steps generally follow that outlined by^9^. To translate relevant genes, we used ANGSD^54^ to create consensus sequences for only those BAM files representing chromosomes with genes of interest. To reduce the effects of post-mortem aDNA damage, we trimmed the first and last five nucleotides from each DNA fragment. We formatted each consensus sequence as a blast nucleotide database. To recover translated protein sequences, we performed a tblastn alignment^55^, with the corresponding *Ceratotherium simum* sequences as queries. Finally, we used ProSplign to recover the spliced alignments, and ultimately, the translated protein sequences^56^.

### Protein identification

Thermo .raw files generated by the mass spectrometer were searched with various softwares using an iterative search strategy to interpret spectra, characterise PTMs, and ultimately, reconstruct protein sequences. For comparison, .raw files from a mediaeval ovicaprine (control) and an early Pleistocene *Stephanorhinus* generated by Cappellini et al. (2019)^9^ were also analysed. Among samples from the Fontana Ranuccio *Stephanorhinus*, only FR4 was analysed.

We primarily employed MaxQuant^57^ for sequence reconstruction and other downstream aspects of data analysis. We performed two initial runs: 1) a more focussed run using the database we modified from Cappellini et al. (2019)^9^, and 2) a broad run using the ‘Theria’-wide database we constructed from publicly-available sequences.

In all runs, an Andromeda threshold of 40 and a delta score of 0 were set for both unmodified and modified peptides. Minimum and maximum peptide lengths were specified as 7 and 25, respectively. The default peptide false discovery rate (FDR) was used (0.01), while protein FDR was increased to 1 to show possible low-abundance proteins. “Unspecific” digestion was specified. No fixed post-translational modifications were set. Several PTMs were set as variable modifications in our initial runs: glutamine and asparagine deamidation, methionine and proline oxidation, N-terminal pyroglutamic acid from glutamic and aspartic acids, phosphorylation of serine, threonine, and tyrosine, and the conversion of arginine to ornithine.

Proteins included in the database of common contaminants provided by MaxQuant (for example proteinaceous laboratory reagents and human skin keratins), as well as reverse sequences, were manually removed and not further examined. In addition, proteins detected in the laboratory blank were also treated as contaminants, and not considered further.

To discover new SAPs and peptide variants not included in our database, we used additional search tools. Peaks v. 7.0 wass used to attempt *de novo* sequencing and homology search was performed using the SPIDER algorithm^58–60^. The open search capabilities of openPFind^61^ and MSFragger^62^ were also used. When possible, the same settings were selected as in the MaxQuant runs.

With our iterative search strategy, we integrated possible sequence variants from the results of our *de novo*, homology searches, and open searches into hypothetical sequences from closely-related taxa, to produce artificial sequences. These artificial sequences were included in a subsequent MaxQuant search, and only incorporated into reconstructed sequences if identified and validated using MaxQuant.

### Sequence reconstruction + filtering

Prior to sequence reconstruction, all non-redundant PSMs were filtered using three criteria to reconstruct only those peptide sequences and amino acid residues that we can confidently assign. Sequences were accepted at two levels, resulting in two different datasets: 1) a minimally-filtered dataset, and 2) a strictly-filtered dataset. This filtering starts with using Basic Local Alignment Search Tool (BLAST)^63^ to determine if peptides match any contaminants, beyond those included in MaxQuant by default, such as soil bacteria and fungi. Next, MS/MS spectra are manually inspected for each PSM to examine ion series coverage. At this stage, peptide sequences are accepted for the strictly-filtered dataset only if each amino acid residue is covered (e.g., at least y-, b-, or a-ion designates the mass of that specific amino acid, plus any identified PTMs) by at least two spectra, following the approach outlined by Coutu et al. (2020)^64^. Additionally, for both strict- and minimally-filtered datasets, poorly-supported spectra are also removed at this stage, and proteins are only submitted for phylogenetic analysis if they are covered by at least two non-overlapping peptides. Finally, under the strict-filtering criteria, BLAST is used again on any trimmed sequences, to remove any that match contaminants.

### Protein damage analysis

Characterization of protein degradation and post-translational modifications roughly follows that of Cappellini et al. (2019)^9^. In addition to the primary run, three additional runs were performed on each rhinocerotid sample, alongside the mediaeval control, to assess several different post-translational modifications: 1) oxidative degradation of tryptophan, and included kynurenine (ΔM = +3.994915), oxolactone (ΔM = +13.979265), tryptophandione (ΔM = +29.974178) as variable modifications, 2) oxidative degradation of histidine (His), including oxohistidine (ΔM = +15.9949), dioxohistidine (ΔM = +31.990), His to hydroxyglutamate (ΔM = +7.979), and His -to aspartic acid (ΔM = -22.032) as variable modifications, and 3) aromatic, including oxidation (WY) (ΔM = +15.9949) and dioxidation (WY) (ΔM = +31.990) as variable modifications. Deamidation (NQ) was also included as a variable modification for each run. After removing potential contaminants and reverse hits, we used spectral counting to assess the extent of each PTM. In the case of arginine, histidine, and tryptophan, we summed the total number of each amino acid residues in each sample, after filtering for reverse hits and potential contaminants, and the total number of each modified amino acid residue in each sample (Figure 2C and Ext. Figure 1BC). Deamidation levels were calculated following the approach described in Mackie *et al*. (2018)^50^ (Figure 2C) and site specific rates were observed using the DeamiDATE algorithm^65^.

### Intra-crystalline protein decomposition analysis

Chiral amino acid analysis was undertaken on CMNF-59632 to evaluate the overall extent of amino acid degradation in the intra-crystalline fraction of the enamel, enabling comparison to previously-analysed specimens^9^, and samples that had been experimentally-heated between 60 and 80°C for up to 17520 hours and to samples heated to 200 - 500 °C for up to 25 min. Enamel chips were drilled using a Dremel ® 4000 (4000-1/45) drill with a diamond wheel point (4.4 mm (7105) by Dremel ®) to remove any dentine which could be identified under a microscope (ZEISS Stemi 305, Axiocam 105 R2). Samples were processed following the methods of Dickinson et al. (2019)^66^. To remove excess powders, enamel chips were washed in deionized water and ethanol (Analytical-grade) before being powdered with an agate pestle and mortar. Powdered samples were weighed into a single plastic microcentrifuge tube and bleached (NaOCl-12%, 50 μL mg^−1^ of enamel) for 72 hours to remove the inter-crystalline amino acids and any contamination. This bleached sample was washed five times with deionized water, and then once with methanol (HPLC-grade), before being left to dry overnight.

The dried bleached sample was then divided into four subsamples: two for replicate analysis of the free amino acids (FAA) and two for replicate analysis for the total hydrolysable amino acids (THAA). The THAA subsamples were dissolved in HCl (7 M, 20 μL mg^−1^, Analytical grade) in a sterile 2 mL glass vial (Wheaton), purged with N_2_ to reduce oxidation and heated at 110 °C for 24 h in an oven (BINDER GmbH series). The acid was then removed by centrifugal evaporation (Christ RVC2-25). Then, THAA and FAA fractions were subjected to a biphasic separation procedure^66,67^ to remove inorganic phosphate from the enamel samples. HCl was added to both FAA (1 M, 25 μL mg^−1^) and THAA (1 M, 20 μL mg^−1^) fractions in separate 0.5 mL plastic microcentrifuge tubes (Eppendorf), and KOH (1 M, 28 μL mg^−1^) was added into the acidified solutions, which then formed mono-phasic cloudy suspensions. Samples were agitated and then samples were centrifuged (13,000 rpm for 10 min, Progen Scientific GenFuge 24D) to form a clear supernatant above a gel. The supernatant was removed, and dried by vacuum centrifugation. The concentration of the intra-crystalline amino acids, and their extent of racemisation (D/L value) were then quantified using RP-HPLC (Agilent 1100 series HPLC fitted with HyperSil C18 base deactivated silica column (5 μm, 250 x 3 mm) and fluorescence detector) following a modified method of Kaufman & Manley (1998)^68^.

For the RP-HPLC analysis, samples were rehydrated with an internal standard solution (L-homo-arginine (0.01 mM), sodium azide (1.5 mM) and HCl (0.01 M)) and run alongside standards and blanks. A tertiary mobile phase system (HPLC-grade acetonitrile:methanol:sodium buffer (21 mM sodium acetate trihydrate, sodium azide,1.3 μM EDTA, pH adjusted to 6.00 ± 0.01 with 10% acetic acid and sodium hydroxide)) was used for analysis. D and L peaks of the following amino acids were separated: aspartic acid and asparagine (Asx); glutamic acid and glutamine (Glx); serine (Ser), alanine (Ala), valine (Val), phenylalanine (Phe), isoleucine (Ile), leucine (Leu), threonine (Thr), arginine (Arg), tyrosine (Tyr) and glycine (Gly). During preparation, asparagine and glutamine undergo rapid irreversible deamination to aspartic acid and glutamic acid respectively^69^ and hence they are reported together as Asx and Glx respectively.

### Phylogenetic Analysis

A time-calibrated phylogenetic tree was inferred with the Bayesian phylogenetic software RevBayes v.1.2.1^70^; https://revbayes.github.io/) under a constant-rate Fossilised Birth Death (FBD) model^71,72^. The dataset consisted of enamel proteome data for 16 perissodactyl species (10 extant and 6 extinct), totalling 7 proteins and 3446 amino acids. Phylogenetic analyses were performed with both the strict-filtered and minimally-filtered sequences for CMNF-59632, to observe any topological differences between the two datasets and assess if filtering is warranted. As no major differences were observed, only the results from the ‘strictly-filtered’ dataset are discussed. The proteome dataset was partitioned by protein. A GTR + I (General Time Reversible + Invariant sites) amino acid substitution model—where stationary frequencies of the 20 amino acids and exchangeability rates among amino acids are free to vary and estimated from data—was applied to each partition. Preliminary unrooted phylogenetic analyses performed on each protein showed evidence for within-protein **Γ**(Gamma)-distributed rate variation only for MMP20, hence **Γ**-distributed rate variation was modelled only for the MMP20 partition. A relaxed clock model with uncorrelated lognormal-distributed rates (UCLN) was applied to allow rate variation across branches. The prior on the average clock rate was set as a loguniform distribution (min=10^-8^, max=10^-2^ substitutions per lineage/million year). The prior on the clock rate standard deviation was set as an exponential distribution with mean equal to 0.587405, corresponding to one order of magnitude of clock rate variation among branches. The FBD tree model allows for placement of extinct species in a phylogenetic tree while simultaneously estimating the rates of speciation, extinction, and fossilisation (sampling of species in the past). The priors on speciation, extinction, and fossilisation parameters were set as uniform distributions bounded between 0 and 10. The sampling probability for extant species was fixed to 0.5882353 (10/17), corresponding to the fraction of extant perissodactyl species included in the analysis, and assuming uniform sampling of extant taxa. The three species of Equidae in the analysis (*Equus caballus*, *E. przewalskii*, *E. asinus*) were constrained as outgroup to other perissodactyls (Tapiridae + Rhinocerotidae). Tip ages of fossil taxa were given a uniform prior distribution ranging from minimum to maximum age of the deposit where each fossil has been found. The prior on the origin age of the tree was set as a uniform distribution with minimum = 54 Ma, corresponding to the oldest fossil that can be unequivocally assigned to crown Perissodactyla (*Cambaylophus vastanensis* from the early Ypresian Cambay Shale^73^), and maximum = 100 Ma, corresponding to the beginning of the Late Cretaceous and a very lax upper boundary on the origin of placental mammals^74^. Additional constraints on node ages based on the fossil record of perissodactyls were set to improve precision of divergence age estimates. Each node calibration was set up as a soft-bounded uniform distribution with normally distributed tails, with 2.5% of the distribution younger than the minimum age (allowing for potential misattribution of the oldest fossil of a clade) and 2.5% of the distribution older than the maximum age. Monophyly was not enforced when setting up these node calibrations. The following age constraints have been applied to five nodes:

1) Node = crown Perissodactyla; soft minimum = 54 Ma, with the same justification as the minimum on the origin age prior; soft maximum = 66 Ma, corresponding to the Cretaceous–Palaeogene boundary, before which no unambiguous crown placental fossils are known. 2) Node = Rhinocerotina (crown rhinoceroses); soft minimum = 22.6 Ma, corresponding to the earliest putative appearance of a crown rhinoceros in the fossil record (*Gaindatherium* cf. *browni* from the Aquitanian upper member of the Chitarwata Formation^75,76^; soft maximum = 44 Ma, corresponding to the minimum age of Rhinocerotidae as supported by fossil and phylogenetic evidence ^25^. 3) Node = Diceroti (*Ceratotherium* + *Diceros*); soft minimum = 5.3 Ma, corresponding to the minimum age of the oldest deposits yielding *Diceros bicornis* fossils (Lothagam and Albertine^77,78^; soft maximum = 7.3 Ma, as in Liu et al. (2021)^25^. 4) Node = *Rhinoceros unicornis* + *Rhinoceros sondaicus*; soft minimum = 1.9 Ma, corresponding to the early Pleistocene appearance of *Rhinoceros unicornis* in the fossil record^79,80^; soft maximum = 5.3 Ma, as in Liu et al. (2021)^25^. 5) Node = *Dicerorhinus* + *Stephanorhinus* + *Coelodonta*; soft minimum = 13 Ma, corresponding to middle Miocene remains of *Dicerorhinus* from the middle Siwaliks of Pakistan^25,81^; soft maximum = 22.6 Ma, corresponding to the oldest crown rhinoceros fossil as in the soft minimum of calibration 2.

The Markov chain Monte Carlo (MCMC) was set up as 4 independent runs, running for 50,000 iterations and sampling every 10, averaging between 262.2 and 279.2 moves per iterations. Convergence between runs was checked by visually inspecting and calculating effective sample sizes (ESSs) of parameter estimates on Tracer v.1.7.2^82^. A maximum *a posteriori* (MAP) tree was calculated to summarise the posterior distribution of trees, with 20% burn-in. In the analysis of the minimally-filtered dataset, one of the 4 runs was discarded from the MAP tree calculation, as it converged only in the last 10% of the MCMC.

## Acknowledgements

This project has received funding from the European Union’s Horizon 2020 research and innovation programme under the Marie Sklodowska-Curie grant agreement (No. 861389 - PUSHH). EC, FZ, IP, KP, FR, JVO, MM and RP are supported by the European Research Council (ERC) under the European Union’s Horizon 2020 research and innovation programme (grant agreement No. 101021361 - BACKWARD). MM was also supported by the Danish National Research Foundation grant PROTEIOS (DNRF128), and MD, KP & CB by the UK NERC (NE/S010211/1 & NE/S00713X/1). DF was supported by the Natural Sciences and Engineering Research Council of Canada (NSERC RGPIN-2018-05305) and a Research Activity Grant through the Canadian Museum of Nature. This work was supported by the Deutsche Forschungsgemeinschaft (DFG) Emmy Noether-Program (Award HO 6201/1-1 to SH) and by the European Union (ERC, MacDrive, GA 101043187). Fieldwork in Canada’s High Arctic was supported by a palaeontology permit from the Government of Nunavut, Department of Culture, Language, Elders and Youth (D.R. Stenton, J. Ross), with the permission of the Qikiqtani Inuit Association, particularly Grise Fiord.

## Author contributions

R.P., R.D.E.M, N.R., D.F., and E.C. designed the study. N.R, D.F. and M.G. conducted fieldwork at the Haughton Crater site. R.D.E.M., N.R., D.F., M.G., R.F., and L.B., provided ancient samples. R.P., M.M, A.B., N.H., I.P., S.L., J.R-M., A.R., F.M., M.R.D., C.B., and G.S. performed data generation and analysed data with support from E.C., S.H., E.W., N.R., R.D.E.M., and D.F.. R.P., M.M, and E.C. wrote the manuscript with contributions from all authors.

## Corresponding authors

Correspondence to Ryan Sinclair Paterson, Danielle Fraser, or Enrico Cappellini.

## Extended Data

**Extended Figure 1).**
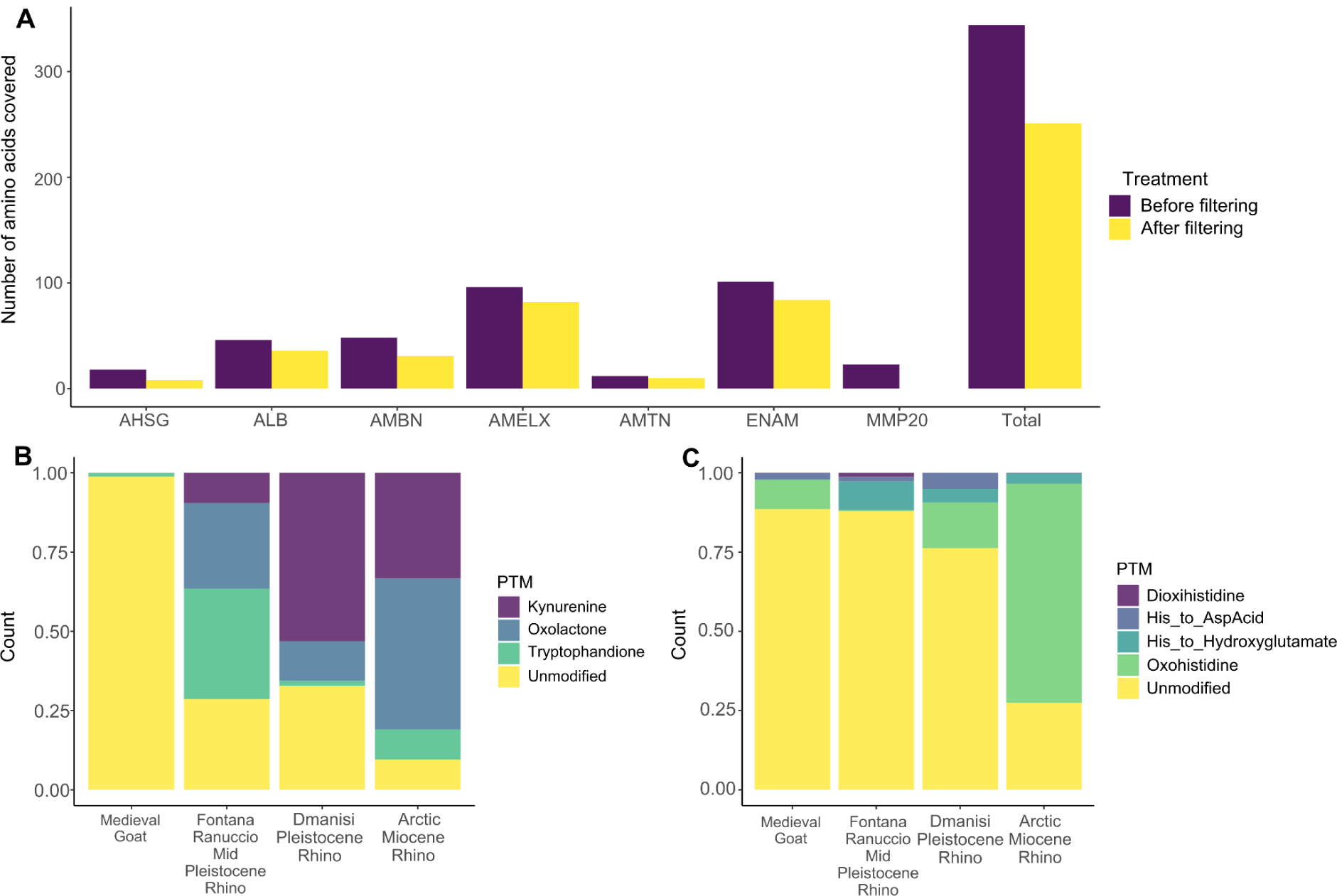
**Proteome preservation in CMNF-59632**. A) Amino acid count for each identified protein, before and after filtering (see Methods), B) PTMs related to oxidative degradation of tryptophan for CMNF-59632, compared to enamel proteomes from other ancient rhinos and a mediaeval ovicaprine, C) PTMs related to oxidative degradation of histidine for same taxon set. The moderate protein preservation in CMNF-59632, indicated by the lower amino acid coverage compared to other ancient enamel proteomes, is further supported by the high incidence of PTMs related to oxidative degradation, compared to other fossil rhinocerotids.

**Extended Figure 2).**
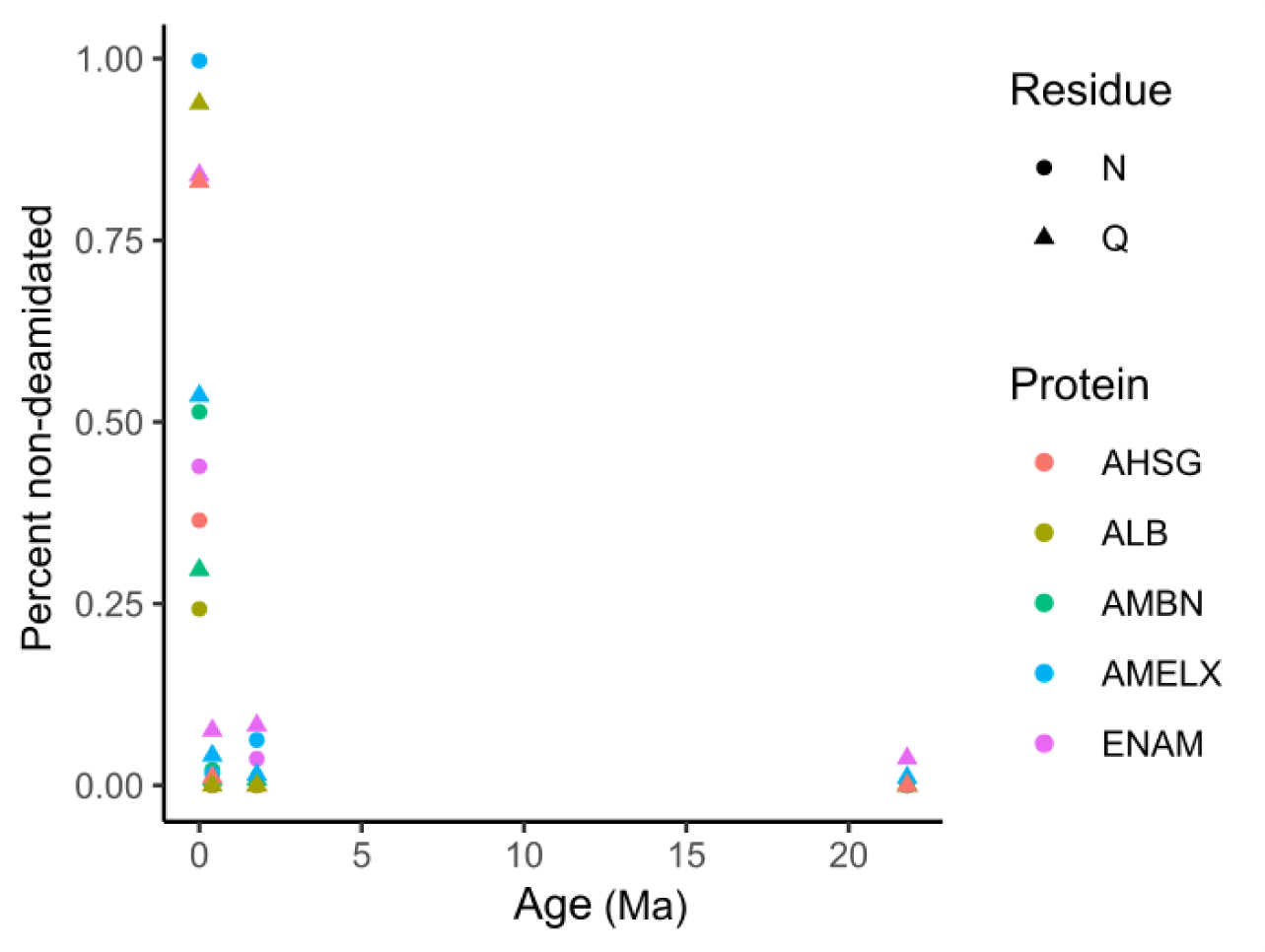
Deamidation rates in fossil rhinocerotid enamel proteomes, plotted against geological age. Data used is CMFN-59632 from Haughton Crater (21.8 Ma), DM.5/157 from Dmanisi (1.77 Ma), CGG 1_023342 from Fontana Ranuccio (0.4 Ma), and a mediaeval control sample (0.005 Ma). While useful for establishing authenticity of an ancient proteome, deamidation rates plateau relatively quickly, so they are not reliable for assessing relative degradative state in ancient proteomes from deep geological timescales.

**Extended Figure 3).**
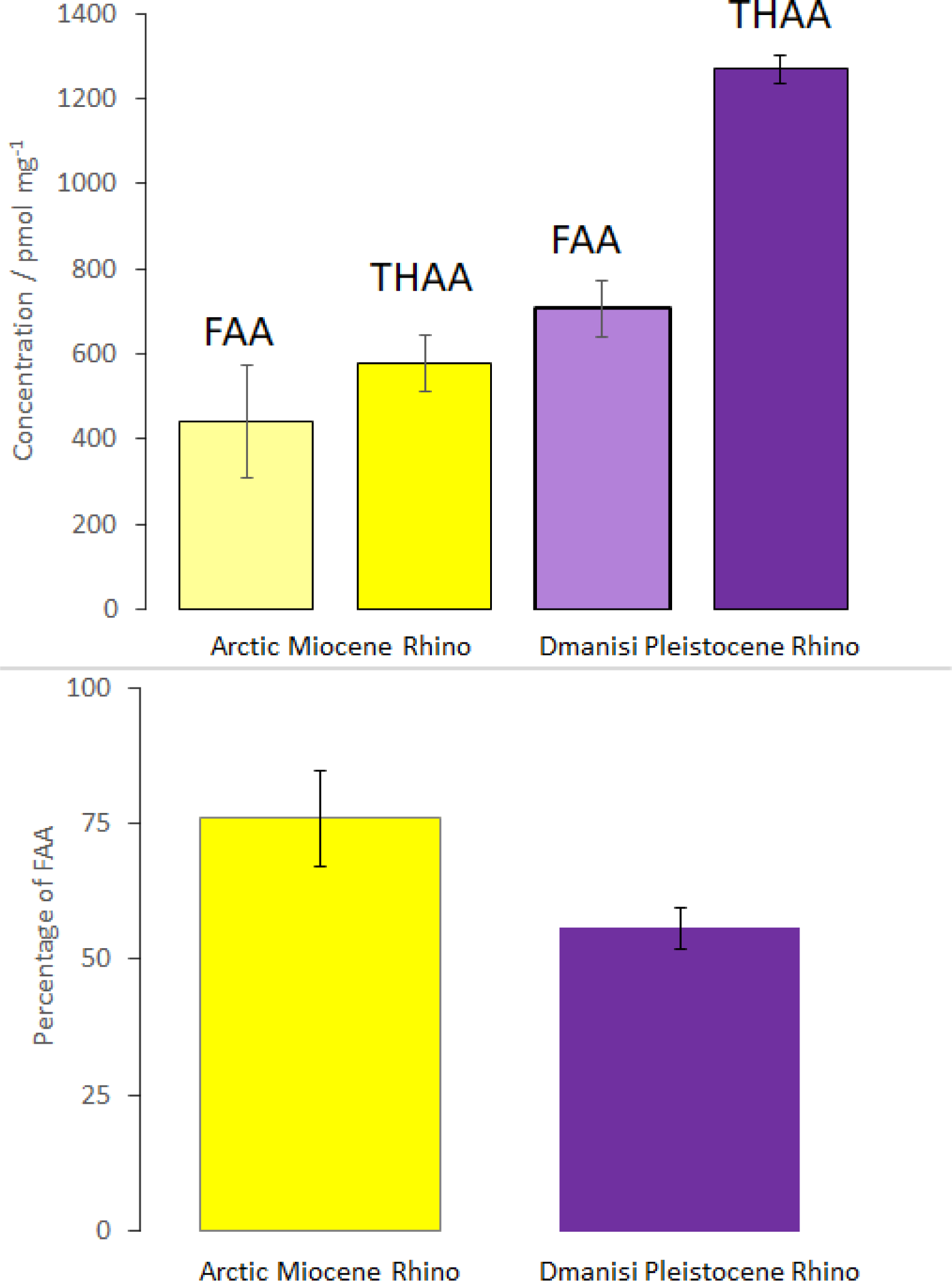
Comparison of FAA & THAA concentration (top) and %FAA (bottom) of the Arctic Miocene rhinocerotid with the Dmanisi Pleistocene *Stephanorhinus^11^*. Error bars represent 1 standard deviation about the mean for preparative replicates. The lower overall concentration, higher %FAA and yet incomplete hydrolysis in the Arctic Miocene rhino is consistent with endogenous peptides in the tooth enamel.

**Extended Figure 4).**
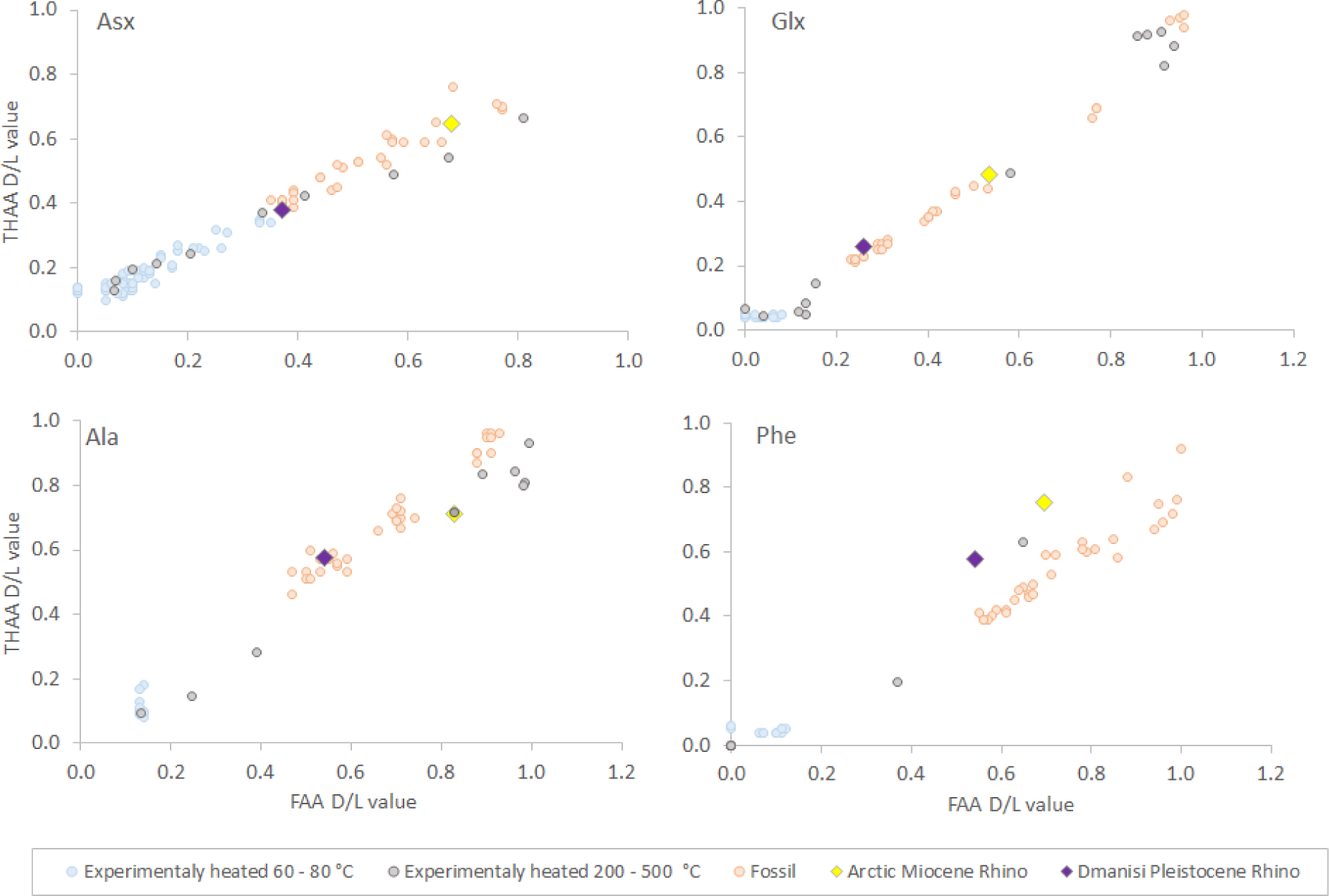
Asx, Glx, Ala and Phe FAA vs THAA D/L values for tooth enamel from Arctic Miocene rhino from Ellesmere Island Canada, and the Dmanisi Pleistocene rhino. A data set consisting of published and unpublished enamel data from other rhino palaeontological and experimental data has been included for comparison. The good correlation between FAA & THAA for the Arctic Miocene rhino (CMNF-59632) sample supports the presence of closed system original peptides and their constituent amino acids in this Miocene sample.

**Extended Figure 5).**
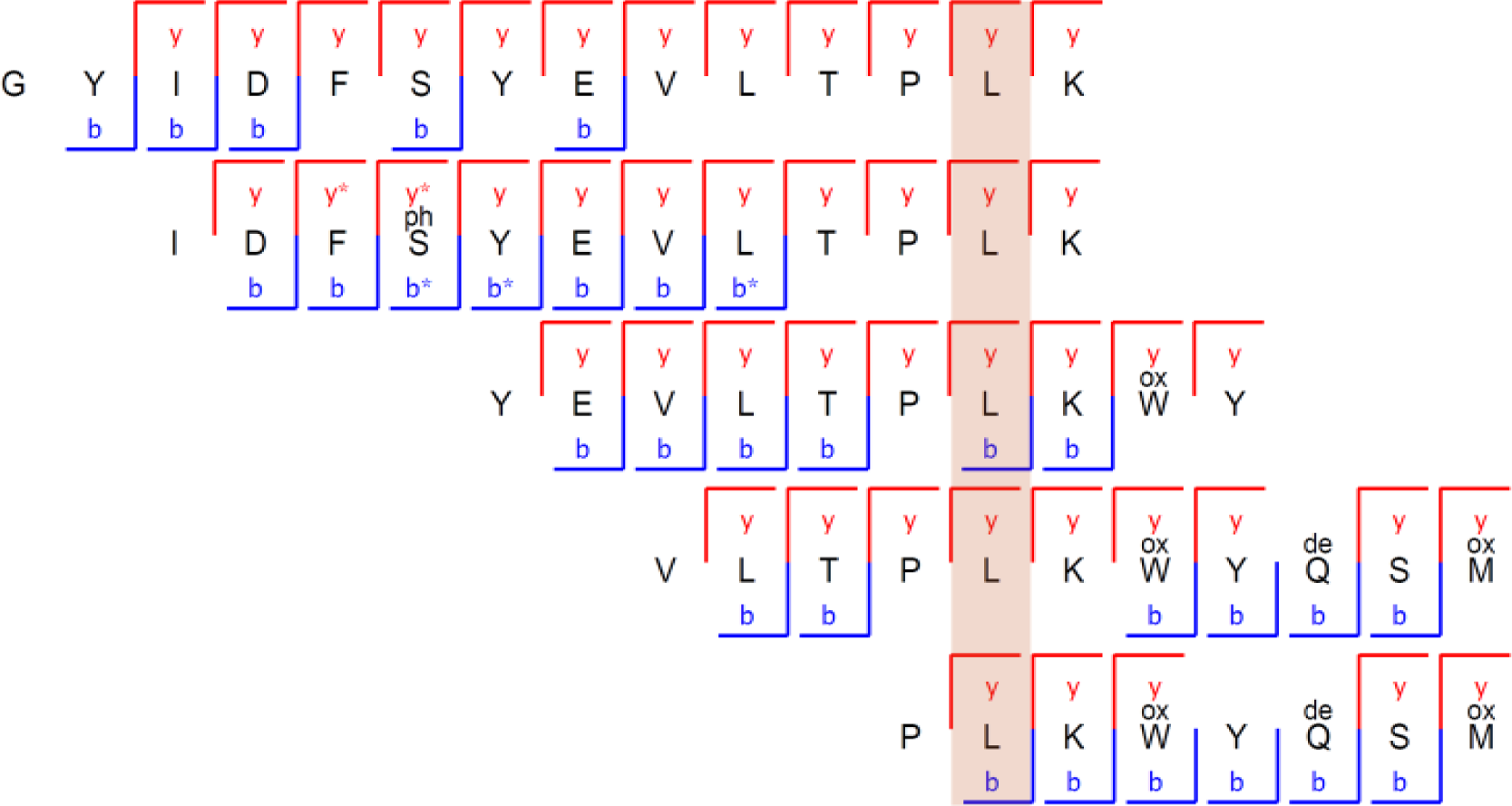
Subset of overlapping peptides supporting a SAP at AMELX position 53. While all other later-diverging rhinocerotids, including *Elasmotherium*, display a valine (V) at position 53 (highlighted in light red; following numbering of reference sequence A0A5F5PLN8), CMNF-59632 displays a leucine (L) (or isoleucine (I)), representing the ancestral condition in Perissodactyla. The peptide sequences depicted here represent just a small portion of the peptide-spectrum matches covering this position. Together, these peptides display high Andromeda scores, extended, in some cases complete, ion series, and the presence of several PTMs supporting their endogeneity (phosphorylation) and ancientness (tryptophan oxidation, glutamine deamidation).

